# DeepPrime2Sec: Deep Learning for Protein Secondary Structure Prediction from the Primary Sequences

**DOI:** 10.1101/705426

**Authors:** Ehsaneddin Asgari, Nina Poerner, Alice C. McHardy, Mohammad R.K. Mofrad

**Affiliations:** Molecular Cell Biomechanics Laboratory, Departments of Bioengineering and Mechanical Engineering, University of California, Berkeley, CA, 94720, USA; Computational Biology of Infection Research, Helmholtz Centre for Infection Research, Brunswick 38124, Germany; Center for Information and Language Processing, Munich 80538, Germany; Molecular Biophysics and Integrated Bioimaging, Lawrence Berkeley National Lab, Berkeley, CA 94720, USA

## Abstract

**Motivation:** Here we investigate deep learning-based prediction of protein secondary structure from the protein primary sequence. We study the function of different features in this task, including one-hot vectors, biophysical features, protein sequence embedding (ProtVec), deep contextualized embedding (known as ELMo), and the Position Specific Scoring Matrix (PSSM). In addition to the role of features, we evaluate various deep learning architectures including the following models/mechanisms and certain combinations: Bidirectional Long Short-Term Memory (BiLSTM), convolutional neural network (CNN), highway connections, attention mechanism, recurrent neural random fields, and gated multi-scale CNN. Our results suggest that PSSM concatenated to one-hot vectors are the most important features for the task of secondary structure prediction.

**Results:** Utilizing the CNN-BiLSTM network, we achieved an accuracy of 69.9% and 70.4% using ensemble top-k models, for 8-class of protein secondary structure on the CB513 dataset, the most challenging dataset for protein secondary structure prediction. Through error analysis on the best performing model, we showed that the misclassification is significantly more common at positions that undergo secondary structure transitions, which is most likely due to the inaccurate assignments of the secondary structure at the boundary regions. Notably, when ignoring amino acids at secondary structure transitions in the evaluation, the accuracy increases to 90.3%. Furthermore, the best performing model mostly mistook similar structures for one another, indicating that the deep learning model inferred high-level information on the secondary structure.

**Availability:** The developed software called DeepPrime2Sec and the used datasets are available at http://llp.berkeley.edu/DeepPrime2Sec.

## 1 Introduction

Proteins are macromolecules that are crucial elements for the structure and function of cells with a wide array of responsibilities including structural support, intra- and inter-cellular transport, catalytic activity, defense against bacteria and viruses, muscle contraction, signaling and regulation. Proteins accomplish their diverse functions in interactions with their environments, which can be other macromolecules (such as proteins, DNA, or RNA), chemical compounds, or factors such as the pH or temperature Clark and Radivojac (2011); Cooper *et al.* (2000). Proteins are polymers of small molecules called amino acids, of which there are 20 different types, represented by the characters {A, C, D, E, F, G, H, I, K, L, M, N, P, Q, R, S, T, V, W, Y}, plus 4 letters indicating ambiguities: *B* instead of {*N* − *or* − *D*}, *J* instead of {*I* − *or* − *L*}, *Z* instead of {*E* − *or* − *Q*}, and *X* as an completely unknown amino acid. Protein sequences can be as large as chains of 10Ks amino acids^1^; meaning that the space of possible protein sequences is very large. Proteins fold to form a particular three-dimensional structure. It has been proven that the protein’s linear sequence can determine their tertiary structures (3D structure) Cooper *et al.* (2000). The functions of proteins are highly tied to their 3D structures. Hence, a protein sequence should theoretically hold enough information determining its function. However, finding a mapping from the protein primary sequence to the structure is one of the open challenges in molecular biology Hunter (1993); Cooper *et al.* (2000). The large gap between the number of known protein sequences (UniProt database contains 116 million protein sequences to date) and the number of known protein 3D structures (Protein Databank contains only 142K entries for protein 3D structures to date) motivates computational methods and in particular machine learning methods predicting structural information from the protein primary sequences. Protein structure can be described at three main levels: **(i) primary structure** referring to a linear sequence of amino acids, **(ii) secondary structure** referring to the structure of the local segments of the protein sequence categorized into 8 of secondary structures, and **(ii) tertiary structure** referring to the 3D structure of protein macromolecules. In this work, we focused on predicting protein secondary structure from the primary sequences.

Protein secondary structure prediction can be viewed as a sequence labeling machine learning task type, i.e. assigning a categorical label *y*_*t*_ ∈ *Y* to each element of a sequence of input elements, *x*_*t*_ ∈ *X*, where *t* indicates the position in the sequence. There exist eight possible secondary structure categories (Q8 labeling) for each amino acid at position *t* in the sequence: the 3-10 helix (G), *α* helix (H), *π* helix (I), turn (T), *β* sheet (E), *β* bridge (B), bend (S), and loop (L). A simpler labeling scheme is Q3, where the categories are divided into three main classes: helix, strand, and loop/coil. Finding the protein secondary structure is an important step toward the understanding of the protein folding and subsequently its structure and function Zhou and Karplus (1999); Ozkan *et al.* (2007). Thus, it can be vital for a variety of protein informatics problem settings, e.g., protein sequence alignment Zhou and Zhou (2005); Deng and Cheng (2011), identification of disease-causing mutations Folkman *et al.* (2015), and protein function annotation Taherzadeh *et al.* (2016).

Secondary structure prediction is one of the primary tasks in protein informatics Yang *et al.* (2016). Traditional protein secondary structure prediction include rule-based Chou and Fasman (1974) methods as well as machine learning approaches using amino acid context-based Finkelstein and Ptitsyn (1971) and evolution/alignment-based representations Zvelebil *et al.* (1987); Hua and Sun (2001). Recently, with the popularity of neural network approaches, similar to many machine learning prediction tasks, different neural network architectures have been proposed for the protein secondary structure prediction from the primary sequences. To the best of our knowledge, all of these architectures have been mainly used on top of a set of fixed features (PSSM and one-hot vector) and mostly on a fixed predictive model in each work. The most challenging dataset for this task has been the 8-way secondary structure prediction using the CullPDB dataset as training and CB513 as the test set Sønderby and Winther (2014). Different deep learning architectures proposed for this task includes: (i) deep Convolutional Generative Stochastic Network, obtaining a test accuracy of 66.4% Zhou and Troyanskaya (2014), (ii) Long Short-Term Memory (LSTM) recurrent neural network, obtaining test accuracy of 67.4% Sønderby and Winther (2014), (iii) Convolutional neural fields achieving a test accuracy of 68.3% Wang *et al.* (2016), (iv) bi-directional LSTM with/without conditional random field (CRF) Johansen *et al.* (2017); Jurtz *et al.* (2017), obtaining a test accuracy of 69.4% and 68.5% respectively, (v) gated Multi-scale convolutional neural network (multi-scale CNN) Zhou *et al.* (2018) with a test accuracy of 69.3% and 70.3%.

The main contributions of the present work are 4-fold, namely (i) we provide a systematic comparison of representations that can be used for the secondary structure prediction task. (ii) We provide a comparison of several important deep learning architectures for this task in order to find the best performing model. (iii) we provide the community with a framework for advancing deep learning techniques in the area of secondary structure prediction, called DeepPrime2Sec (iv) We perform a detailed analysis of the location and classes of errors.

We first implement one of the state-of-the-art neural architecture for sequence labeling tasks in different domains (e.g., proteomics Jurtz *et al.* (2017), natural language processing Lample *et al.* (2016); Taslimipoor and Rohanian (2018); Rohanian *et al.* (2019)). Subsequently, we investigate the role of different features in the performance of protein secondary structure prediction. In particular, we utilize five sets of features and their combinations: (i) one-hot vector representation, (ii) biophysical scores of amino acids, (iii) amino-acid protein vectors, (iv) recently introduced contextualized embeddings, and (v) Position-Specific Scoring Matrix (PSSM). Secondly, for the best feature set in the preceding step, we investigate the performance of different deep learning architectures for the task of secondary structure, including convolutional-recurrent neural network (with and without conditional random field layer), highway connections, attention mechanism, and multi-scale CNN models. Our results confirm that PSSM is the most informative feature for the protein secondary structure, and other features only result in slight improvements when they are combined with PSSM. Error analysis indicates that errors are mostly occurring at positions were transitions between successive secondary structure elements occur. We provide the Prime2Sec code for further investigations.

## 2 Materials and Methods

### 2.1 Datasets

#### Secondary structure prediction dataset

Several benchmark datasets exist in the literature of protein secondary structure prediction. The most challenging dataset based on the maximum achieved accuracy using machine learning approaches is the CullPDB dataset, which consists of 5,534 protein sequences (CullPDB-train) for training and 513 non-redundant sequences (CB513) for test purpose Zhou and Troyanskaya (2014), in which sequences with more than 25% sequence identity to training data entries were removed from the validation set.

The label for each position is selected from the Q8 scheme (explained in § 1). We used as performance metric accuracy, which is the most common metric for the evaluation of protein secondary structure predictors over the filtered CB513. Accuracy can be defined as the ratio of correctly predicted secondary structures in amino acid level.

#### UniRef50 dataset for ELMo training

We use UniRef50 collection as the training data to learn the contextualized embeddings. The primary purpose of using this dataset instead of the whole Swiss-Prot or UniProt has been to avoid having redundant sequences in the test set. We use 90% of sequences for training and 10% for test purpose.

#### Swiss-Prot for ProtVec training

As we proposed in Asgari and Mofrad (2015); Asgari *et al.* (2019) for the training of ProtVec embedding, we use the whole Swiss-Prot dataset containing 600Kprotein sequences.

### 2.2 Approach

First, to investigate the effect of different representations on secondary structure prediction performance, we fix the predictive model and investigate the accuracy under representation changes. For this purpose, we re-implement the state-of-the-art architecture for sequence labeling tasks, i.e., convolutional bidirectional LSTM model Johansen *et al.* (2017). The general architecture used for secondary structure in this paper is illustrated in 1 (a). Secondly, we examine the impact of several deep learning architectures on top of the best feature set obtained in the first step. Thirdly, we create an ensemble predictor on the best performing models. Finally, we present an error analysis of the misclassification locations and confusing secondary structure categories. The experiment steps are detailed as follows.

#### Investigation on the contribution of features in protein secondary structure prediction

We experiment on five sets of protein features to understand what are essential features for the task of protein secondary structure prediction. Although in 1999, PSSM was reported as an important feature to the secondary structure prediction Jones (1999), this was still unclear whether recently introduced distributed representations can outperform PSSM in such a task. For a systematic comparison, the features detailed as follows are used:

- **One-hot vector representation (length: 21)**: vector representation indicating which amino acid exists at each specific position, where each index in the vector indicates the presence or absence of that amino acid.
- **ProtVec embedding (length: 50)**: representation trained using Skip-gram neural network on protein amino acid sequencesAsgari and Mofrad (2015); Asgari *et al.* (2019), detailed in *§*2.2. The only difference would be character-level training instead of n-gram based training.
- **Contextualized embedding (length: 300)**: we use the contextualized embedding of the amino acids trained in the course of language modeling Peters *et al.* (2018), known as ELMo, as a new feature for the secondary structure task. Contextualized embedding is the concatenation of the hidden states of a deep bidirectional language model. The main difference between ProtVec embedding and ELMO embedding is that the ProtVec embedding for a given amino acid or amino acid k-mer is fixed and the representation would be the same in different sequences. However, the contextualized embedding, as it is clear from its name, is an embedding of word changing based on its context. We train ELMo embedding of amino acids using UniRef50 dataset in the dimension size of 300.
- **Position Specific Scoring Matrix (PSSM) features (length: 21)**: PSSM is amino acid substitution scores calculated on protein multiple sequence alignment of homolog sequences for each given position in the protein sequence.
- **Biophysical features (length: 16)** For each amino acid we create a normalized vector of their biophysical properties, e.g., flexibility Vihinen *et al.* (1994), instability Guruprasad *et al.* (1990), surface accessibility Emini *et al.* (1985), kd-hydrophobicity Kyte and Doolittle (1982), hydrophilicity Hopp and Woods (1981), and etc.

##### Feature combinations

We start from using single features and then greedily use the best combinations. Since the PSSM features have a significant contribution in improving the accuracy, we select then for every combination of length two feature sets and more. We perform an extensive parameter tuning of the network hyper-parameters (LSTM size, CNN filter sizes, etc.) for each feature combination.

##### Data augmentation for the best set of features

as later will the reader will see in the results, the combinations of one-hot and PSSM lead to the best performance on the test set. Since PSSM has information about the possible substitution of amino acids in the homolog sequences and homologous sequences are likely to have similar functions and structures, we come up with the idea of generating more data points using PSSM feature (based on most possible mutations) and test it as a data-augmentation policy. This way, we produce ten possible samples from each training instance by changing the amino acid (or more accurately keeping PSSM the same and change the one-hot by sampling from PSSM vector).

#### Deep learning models for protein secondary structure prediction

In addition to investigation on the relevant features, we use different deep learning architectures on top of the selected feature in §2.2 and examine the role of architecture for the protein secondary structure prediction. In a general notation, each sequence labeling task can be viewed as assigning a sequence of labels **Y** = (*y*_1_, *y*_2_, …, *y*_*T*_) to the chain of elements in a given input sequence **X**= (*x*_1_, *x*_2_, …, *x*_*T*_). Using neural architectures, we look for **Y**\* maximizing *P*_θ_(**Y**|**X**). *P*_θ_ is parameterized using a softmax function:

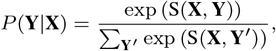

where *S* is a score function computed through a neural network relating the input **X** to the output **Y**. The difference between different models is in the neural architecture parametrizing the S. We study the use of various deep learning architectures to produce *S* described as follows.

a. **CNN-BiLSTM Model**: As the CNN-BiLSTM model is illustrated in Figure 1 (a), firstly convolutional filters of different window sizes are applied on the input features, creating feature maps of different neighborhoods. Then the feature maps are concatenated. The resulting vector encodes the representation of different context sizes around each amino acid in the protein sequence. Batch normalization is used to increase the stability of training Ioffe and Szegedy (2015). Subsequently, a fully-connected neural network projects the result into a dense vector. In order to avoid over-fitting, dropout is used Srivastava *et al.* (2014). Up to this point we encoded a sequence of amino acid features vectors **X** = (*x*_1_, *x*_2_, …, *x*_*T*_) into a dense vector **V** = (**v**_1_, **v**_2_, …, **v**_*T*_) using a convolutional and feedforward neural network, which is enriched on local information at each position *t*. However, *v*_*t*_ does not encode global information about the sequence. A long Short-term Memory Network (LSTM) Hochreiter and Schmidhuber (1997), which is designed to capture long-range dependencies, encodes the sequence information. The LSTM creates an encoding 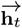 of the left context of the protein at position *t*. Since both left and right contexts can be crucial for the global structure of proteins, we use a bi-directional LSTM (or shortly BiLSTM network). The Bi-LSTM encodes each position into a representation of left and right contexts 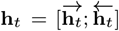. Utilizing feedforward layers (with dropout) on top of **h**_*t*_, we create a vector Φ with a length of the number of possible target labels (|*Y*| = 8). Φ_*t*_ can be regarded as *S*(*x*_*t*_, *y*). Applying a softmax function on the score vector *S*, the label with the maximum value in *S* is chosen.
b. **CNN-BiLSTM with Highway Connections:** the importance of PSSM features in the secondary structure prediction motivates a more direct application of this representation in the last layer as a highway connection He *et al.* (2016). We concatenated the output of BiLSTM **h**_*t*_ with the highway connection to batch-normalized PSSM feature.
c. **CNN-BiLSTM with Conditional Random Field (CRF) layer:** in some natural language processing sequence labeling tasks where there is a complex dependency between neighbor labels, e.g., named entity recognition tasks, using a conditional random field layer on top of the recurrent neural network enhances the prediction power by adding constraints to the final predicted labels Lample *et al.* (2016). A similar idea has been applied in Johansen *et al.* (2017) to the protein secondary structure prediction. In this model, instead of parametrizing the *S* only with the output of the Bi-LSTM model, we use a CRF layer to consider the transition probability between neighboring labels as well:

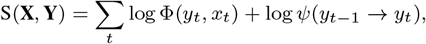

where Φ comes from the output of the BiLSTM and the subsequent feedforward layer and the *Ψ*(*y*_*t*−1_ → *y*_*t*_) is the potential function in the CRF model.
d. **CNN-BiLSTM with Attention layer:** another approach for defining *S* is to write it as the weighted average of all LSTM hidden states. This way, we would allow the model to benefit from the weighted long-term dependencies (global context of the current amino acid). The modification is shown in Figure 1 (d):

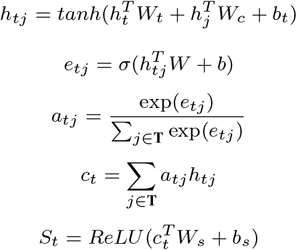

where *h*_*t,j*_ is the contribution of each previous and future LSTM steps in the current state, and *a*_*t,j*_ is the normalized step’s contributions (attention weight), which is used to weight the LSTM *h*′_*x*_*s*. The final encoding for the time *t* would be *c*_*t*_ instead of *h*_*t*_. Finally a non-linear transformation of *c*_*t*_ creates a vector in size of the targets (8 classes) representing the score function *S*.
e. **CNN:** An alternative architecture for this task would be the pure use of the convolutional neural network on the sequence axis (the order in the sequence) to capture local information for prediction on the secondary structure.
f. **Multiscale-CNN with a highway connection:** we implemented one of the state-of-the-art architectures proposed in Zhou *et al.* (2018) as part of *DeepPrime2Sec*, which uses a gated version of stacked CNNs. The ultimate output of each convolutional layer would be a gated version of its input and the convolutional result as depicted in Figure 1 (f):

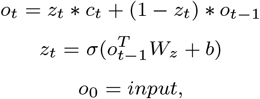

where *c*_*t*_ is the result of the *t*^th^ convolutional layer, *o*_*t*_ is the out of gating between the previous and the current convolutions. In the first layer, *O*_0_ is the same as the input layer.

**Fig. 1:**
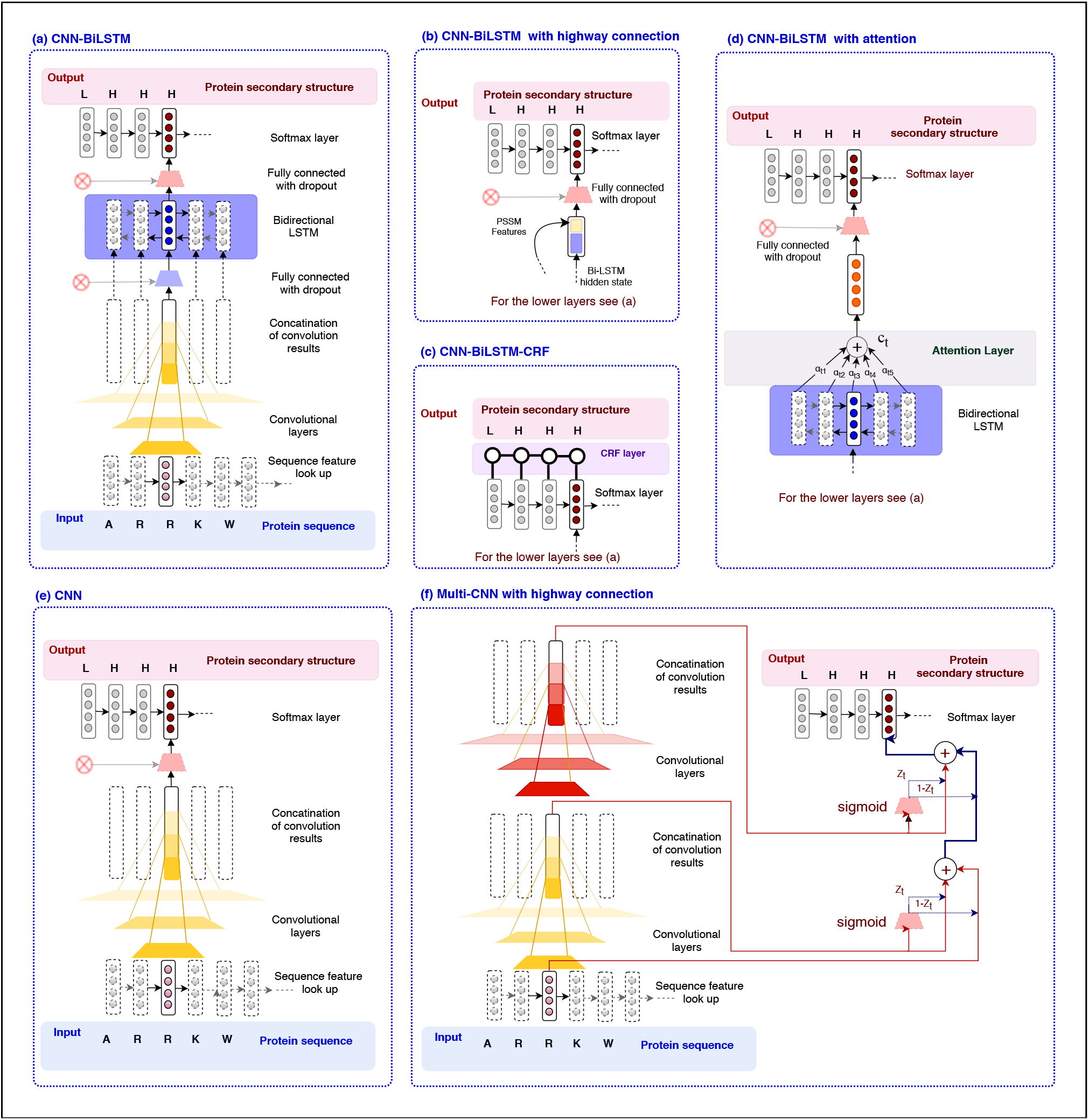
Different deep learning architectures implemented and evaluated in DeepPrime2Sec are provided as follows: **(a)**CNN-BiLSTM, The central neural architecture we used for protein secondary structure; **(b)**The modification of the model (a) with adding highway connection to the CNN-BiLSTM for a more direct use of PSSM features; **(c)**The modification of the model (a) by adding a CRF layer to consider label consistency in the prediction; **(d)**The modification of the model (a) by adding an attention layer to the output of LSTM for considering a more global context; **(e)**Solely using convolutional layer **(f)**Multiscale convolutional neural network benefiting from a gating mechanism (as proposed in Zhou *et al.* (2018)).

**Ensemble of the best models:** similar to previous work, to improve this performance further using “Wisdom of Crowds principle”, we produced an ensemble classifier on the top-k classifiers (k=5,10,20,50,100) and predict on the test set by voting.

## 3 Results

### 3.1 Results on the role of features in secondary structure prediction

#### Single features

The performance of protein secondary structure prediction using different combinations of features is provided in Table 1. When we used single feature sets, PSSM features performed substantially better than other features. Even recent deep learning-based representations were far behind. PSSM is amino acid substitution scores calculated on protein multiple sequence alignment of homolog sequences for each given position in the protein sequence. This result confirms that evolution has very important information for defining the protein structure. One-hot vector encoding performed similar to other amino-acid embedding approaches (ELMo, ProtVec, and biophysical features). The reason behind can be that the one-hot vectors and protein embeddings are acting complementary to each-other; one-hot vector representation increases the precision about the specific residue at position *t*, while embedding blurs this information by providing information about the context of this amino acid. Among the embedding methods, ELMo embedding worked marginally better than other approaches, as it has information about long-term past and future of the sequence, i.e., having information on a more the global context) in comparison to ProtVec, which only includes information about a local context within a certain context size.

**Table 1.** Protein secondary structure prediction results using (i) different feature types, (ii) their combinations, (iii) extensively tuned hyperparameters for the best feature sets, (iv) the data augmentation are presented.

| Representation | Convolution filters | LSTM input/hidden/output | CB513 Q8 accuracy |
| --- | --- | --- | --- |
| (i) Single Features |  |  |  |
| Biophysical features (16D) | [3, 5, 7], 16x | 400 - 800 - 400 | 0.573 |
| ELMo embedding (300D) | [3, 5, 7], 128x | 400 - 800 - 400 | 0.577 |
| ProtVec embedding (50D) | [3, 5, 7], 128x | 400 - 800 - 400 | 0.573 |
| PSSM (21D) | [3, 5, 7], 128x | 400 - 800 - 400 | 0.694 |
| One-hot (21D) | [3, 5, 7], 16x | 200 - 400 - 200 | 0.575 |
| (ii) Combinations of top features |  |  |  |
| Biophysical features & PSSM | [3, 5, 7], 256x | 1000 - 1000 - 1000 | 0.697 |
| ELMo & PSSM | [3, 5, 7], 256x | 400 - 800 - 400 | 0.692 |
| ProtVec embedding & PSSM | [3, 5, 7], 256x | 1000 - 1000 - 1000 | 0.696 |
| One-hot & PSSM | [3, 5, 7], 256x | 1000 - 1000 - 1000 | **0.698** |
| Biophysical features & ELMo & PSSM | [3, 5, 7], 256x | 400 - 800 - 400 | 0.692 |
| Biophysical features & ProtVec embedding & PSSM | [3, 5, 7], 256x | 500 - 1000 - 500 | 0.694 |
| Biophysical features & one-hot & PSSM | [3, 5, 7], 256x | 2000 - 1000 - 2000 | **0.699** |
| ELMo & ProtVec embedding & PSSM | [3, 5, 7], 256x | 400 - 800 - 400 | 0.692 |
| ELMo & one-hot & PSSM | [3, 5, 7], 256x | 1000 - 1000 - 1000 | 0.692 |
| ProtVec embedding & one-hot & PSSM | [3, 5, 7], 128x | 400 - 800 - 400 | 0.694 |
| Biophysical features & ELMo & ProtVec embedding & PSSM | [3, 5, 7], 256x | 1000 - 1000 - 1000 | 0.69 |
| Biophysical features & ELMo & one-hot & PSSM | [3, 5, 7], 256x | 500 - 1000 - 500 | 0.693 |
| Biophysical features & ProtVec embedding & one-hot & PSSM | [3, 5, 7], 256x | 500 - 1000 - 500 | 0.694 |
| ELMo & ProtVec embedding & one-hot & PSSM | [3, 5, 7], 256x | 500 - 1000 - 500 | 0.692 |
| Biophysical features & ELMo & ProtVec embedding & one-hot & PSSM | [3, 5, 7], 256x | 1000 - 1000 - 1000 | 0.692 |
| (iii) Parameter tuning for the selected features |  |  |  |
| One-hot & PSSM | [3, 5, 7, 11, 21], 256x | 1000 - 1000 - 1000 | **0.699** |
| (iv) Data augmentation for the selected parameter setting |  |  |  |
| Augmented one-hot & PSSM in training | [3, 5, 7, 11, 21], 256x | 1000 - 1000 - 1000 | 0.692 |
| Augmented one-hot & PSSM in testing | [3, 5, 7, 11, 21], 256x | 1000 - 1000 - 1000 | 0.68 |

#### Combination of features

The second set of rows in Table 1 shows the combinations of features with PSSM (the best performing feature) starting from combinations of two types up to five feature types. We also optimized the hyperparameters (network sizes and parameters). The combination of one-hot and PSSM turned out to be the most effective representation, while using combination of features did not substantially improve the accuracy further. A reason could be that the PSSM implicitly already includes information about the biophysical and contextual features, at least as much as needed for the protein secondary structure prediction, more than other embedding approaches. Further tuning of the convolution window sizes resulted in the accuracy of 69.9% (Table 1).

#### Data augmentation

Although augmenting the dataset by keeping the PSSM and generating more one-hots made the training dataset 10x larger; the performance could not be improved further 1. Next, we generate ten instances from each test instance and used the best model to predict 10 secondary structures for each position, based on the augmented test set and take the majority vote. This idea also did not boost performance. We conclude that the most important feature to this task is the PSSM, and we need to find a way to perform augmentation on this feature for further performance improvements.

### 3.2 Results on comparison of deep learning architectures in protein secondary structure prediction

The results on secondary structure prediction for different deep learning architectures using the best feature set (i.e., the combination of PSSM and one-hot representation, see §3.1) is provided in Table 2. Using only BiLSTM and only CNN, we could achieve accuracies of 67.1% and 68.1%, respectively. Combination of CNN and BiLSTM led to the best observed accuracy of 69.9%. Adding the CRF layer, attention layer, and the highway connection did not further improve the performance. From this, we may conclude that the LSTM hidden states stored sufficient information, ensuring to provide a logical sequence of labels. The previously reported performances on the *CB513* are also provided in Table 2. The CNN-BiLSTM architecture outperformed all, except for the *CNNH_PSS* model. Based on the provided implementation and descriptions in the Zhou *et al.* (2018), we attempted to reproduce the architecture and the results in the *DeepPrime2Sec*. However, using this architecture, we could not obtain better accuracy than 68.0%.

**Table 2.**
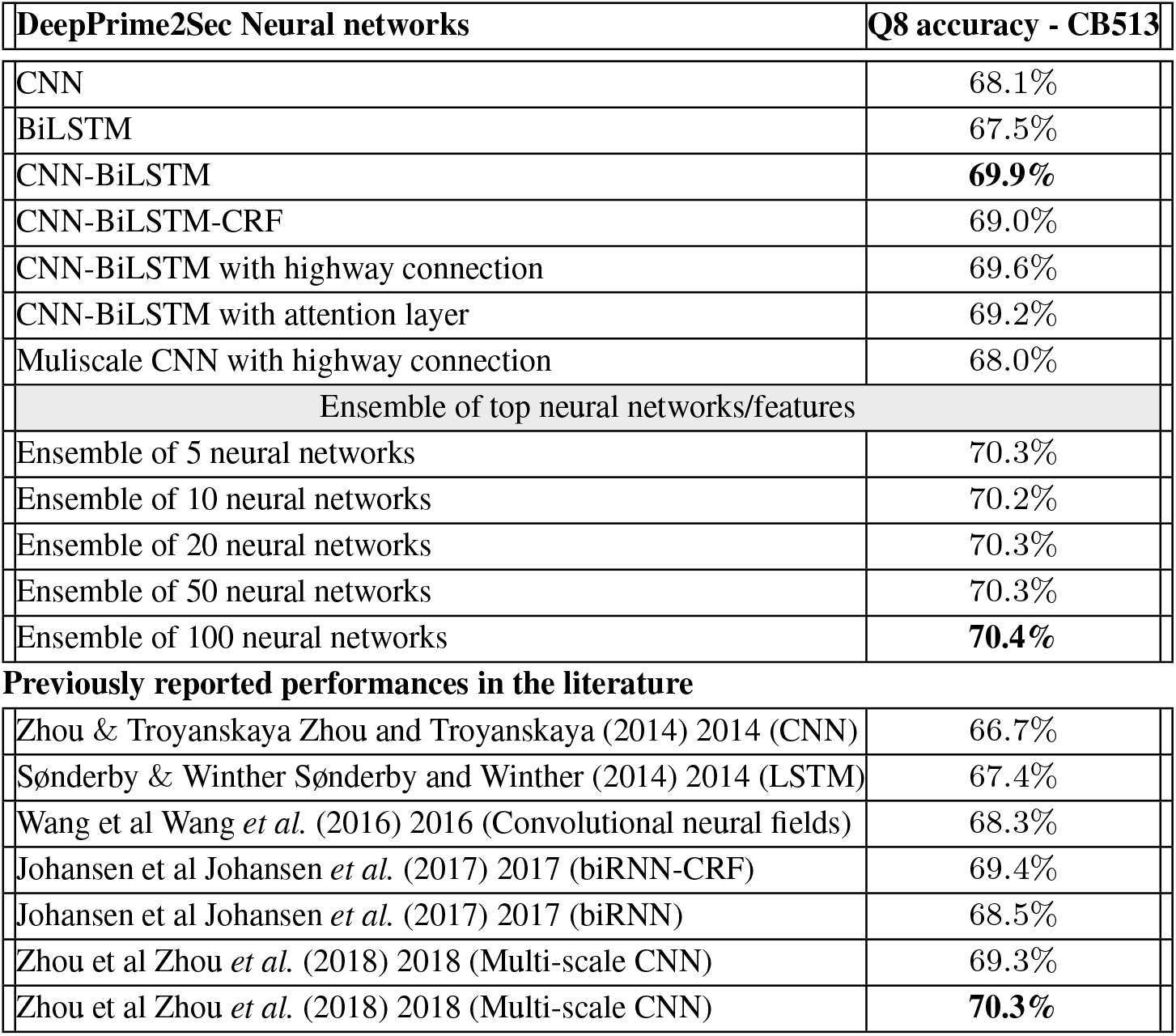
Protein secondary structure prediction results on different deep learning architectures implemented in DeepPrime2Sec, on top of the combination of PSSM and one-hot representation and the ensemble of their top-k models are shown and compared to the state-of-the-art approaches on the CB513 test set.

| DeepPrime2Sec Neural networks | Q8 accuracy - CB513 |
| --- | --- |
| CNN | 68.1% |
| BiLSTM | 67.5% |
| CNN-BiLSTM | **69.9%** |
| CNN-BiLSTM-CRF | 69.0% |
| CNN-BiLSTM with highway connection | 69.6% |
| CNN-BiLSTM with attention layer | 69.2% |
| Multiscale CNN with highway connection | 68.0% |
| Ensemble of top neural networks/features |  |
| Ensemble of 5 neural networks | 70.3% |
| Ensemble of 10 neural networks | 70.2% |
| Ensemble of 20 neural networks | 70.3% |
| Ensemble of 50 neural networks | 70.3% |
| Ensemble of 100 neural networks | **70.4%** |
| Previously reported performances in the literature |  |
| Zhou & Troyanskaya Zhou and Troyanskaya (2014) 2014 (CNN) | 66.7% |
| Sønderby & Winther Sønderby and Winther (2014) 2014 (LSTM) | 67.4% |
| Wang et al Wang et al. (2016) 2016 (Convolutional neural fields) | 68.3% |
| Johansen et al Johansen et al. (2017) 2017 (biRNN-CRF) | 69.4% |
| Johansen et al Johansen et al. (2017) 2017 (biRNN) | 68.5% |
| Zhou et al Zhou et al. (2018) 2018 (Multi-scale CNN) | 69.3% |
| Zhou et al Zhou et al. (2018) 2018 (Multi-scale CNN) | **70.3%** |

#### Ensemble predictor

To further improve this performance, we produced an ensemble classifier on top-k classifiers (k=5,10,20,50,100) resulting in the accuracy of 70.4 (Table 1) outperforming the 70.3 the state-of-the-art performance Zhou *et al.* (2018).

#### Location analysis of misclassified amino acids

Predicting the secondary structure of amino acids at the transition points (transition between two distinct secondary structure) can be tricky. We had the hypothesis that transition positions may have a larger chance of misclassification, as even the ground-truth quality can be lower in these states Yang *et al.* (2016). To study the effect of amino acid position in the misclassifications, we performed statistical tests to determine whether the misclassification event is dependent or independent of locating to the boundaries of the secondary structures in the *CB*513 target labels. We performed both *χ*^2^ and log-likelihood ratio (i.e., the *G*-test) tests and found that the misclassification highly depends on being at the transition location (the p-values on both *χ*^2^ - and *G*- tests were ≈ 0). The underlying contingency table is shown in Table 3. Surprisingly, if we omit amino-acids at the borders from the evaluation, the Q8 accuracy would increase to 90.3%.

**Table 3.**
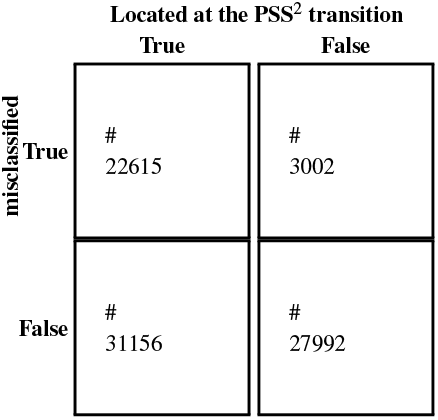
Contingency table for location analysis of the misclassified amino acids

|  |  | Located at the PSS<sup>2</sup> transition |  |
| --- | --- | --- | --- |
|  |  | True | False |
| misclassified | True | #<br>22615 | #<br>3002 |
|  | False | #<br>31156 | #<br>27992 |

#### A categorical analysis of the misclassified amino acids

The confusion matrix and *l*1 normalized confusion matrix for the best performing model using the combination of PSSM and one-hot vector in CNN-BiLSTM architecture are presented in Figure 2 (a) and (b) respectively. Since the protein secondary structure problem setting is relatively imbalanced, the normalized confusion matrix in Figure 2 (b) can more clearly show which classes are relatively confused with each other. The most common secondary structure classes of the alpha helix (H), loop (L), beta sheet (E) were predicted accurately. The classes of bend (B), turn (T) were considerably confused with loop (L), which makes sense, as these classes are similar to each-other and loop (L) is the most frequent one among them. As expected based on structural similarities, bend (S), turn (T), and loop (L) as well as 3-10 helix (G) and alpha helix (H) were also highly confused.

**Fig. 2:**
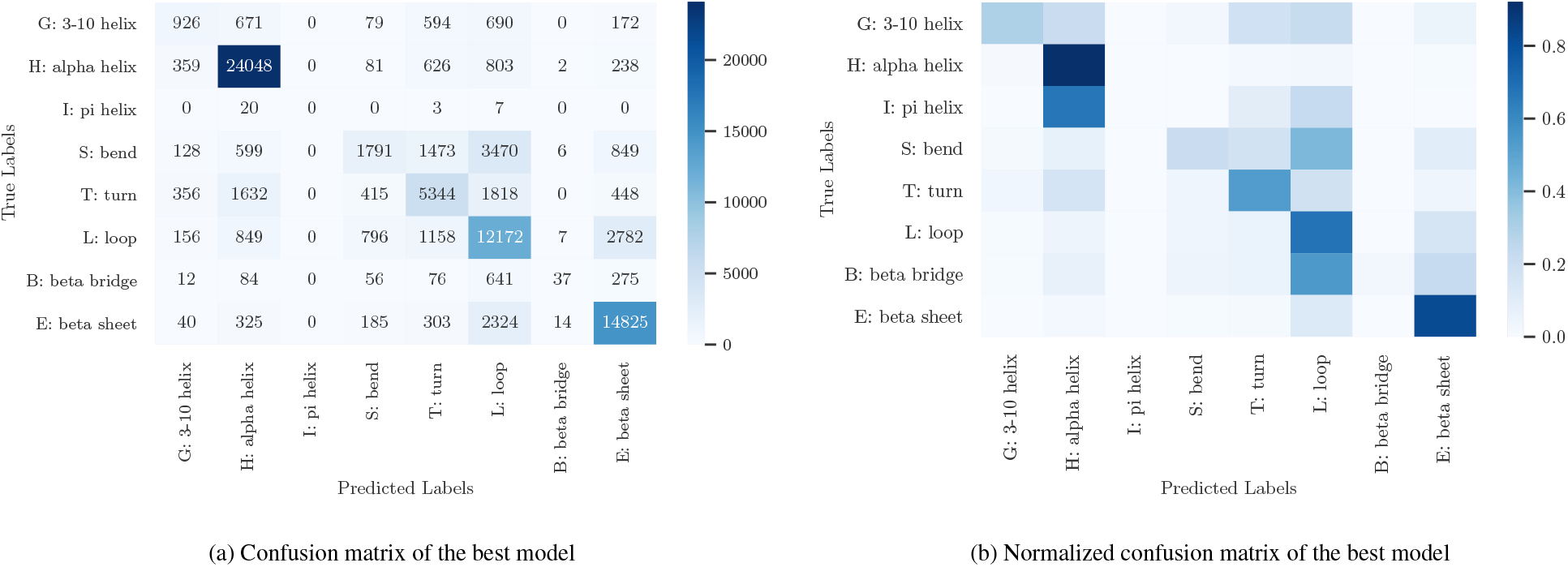
The confusion matrices of the best performing predictor for protein secondary structure prediction on the test data (on CB513) are provided. Figure (a) shows the standard confusion matrix. However, since the secondary structures are not balanced, for better visualization class confusions, we have *l*1 normalized the rows in figure (b).

## 4 Discussions and Conclusion

We studied the machine learning-based protein secondary structure prediction approaches from the protein primary sequence. We focused on finding an optimal representation and deep learning predictive model for this task. The most challenging dataset for this task to-date is Q8 (8 classes) on CullPDB/CB513 dataset, where the dissimilarity of training and test set is ensured. We investigated (i) different protein sequence representations including one-hot vectors, biophysical features, protein sequence embedding (ProtVec), deep amino acid contextualized embedding (ELMo), and the Position Specific Scoring Matrix (PSSM), (ii) different deep-learning architectures including convolutional neural networks (CNN), recurrent neural networks (in particular Bi-LSTM), use of highway connection, attention mechanism, and multi-scale CNN Zhou *et al.* (2018). We showed that PSSM and its combination with one-hot vectors achieve the best performance in protein secondary structure prediction. The best performing model was the CNN-BiLSTM architecture, which captures both local and global sequence features essential for proteins secondary structure. We explored data augmentation of one-hot vector based on the PSSM, which was not successful. A future direction could be exploring possible data augmentation schemes of PSSM features.

*DeepPrime2Sec* provides the community with a deep learning tool specialized for the protein secondary structure prediction covering different architectures. The BiLSTM-CRF architecture performs competitive to the other existing approach in the literature, and the ensemble of the best performing model in Prime2Sec marginally outperforms the existing methods.

In addition, we performed error analysis on the most accurate model based on the location of misclassified amino acids as well as the confusion matrix analysis. Strikingly, misclassified secondary structures were significantly correlated with locating at the structural transitions. Such a correlation is most likely due to the inaccurate assignment of the secondary structure at the boundaries in ground-truth Yang *et al.* (2016). By ignoring the boundary amino acids from the evaluation, the Q8 accuracy would increase for an extra 20%, i.e., 90.3%. Analysis of the confusion matrix furthermore indicates that similar secondary structures are highly confusing (helices: H and G as well as unstructured regions: S, T, and L) showing that the model can learn high-level information about the secondary structures. Even if the exact secondary structure was not predicted, the predicted structure is similar to the target structure.

## Acknowledgements

Fruitful discussions with Omid Rohanian, Hinrich Schütze, Vikrum Singh Nibber are gratefully acknowledged.

## Footnotes

1 based on sequence lengths on UniProt

